# Neuronal excitability and parameter variability in the Hodgkin-Huxley model

**DOI:** 10.64898/2025.12.30.696982

**Authors:** Alon Korngreen

## Abstract

Biophysically detailed neuron models are often built as a one-way pipeline in which voltage-clamp data are reduced to a single set of best-fit channel parameters, which are then combined into a deterministic spiking model. This practice discards experimentally observed variability and obscures the mechanisms by which robustness and degeneracy arise in excitable systems. Here, we reintroduce parameter variability into the Hodgkin-Huxley model and embed uncertainty and global sensitivity analysis into model construction. We digitized sodium and potassium rate constant data from the original Hodgkin and Huxley figures and used bootstrap resampling to estimate the variability of the voltage-dependent kinetic parameters. We then propagated these uncertainty estimates through a spatially extended squid axon cable model using large-scale Monte Carlo simulations. At the channel level, first-order Sobol sensitivity indices revealed that all kinetic parameters contribute to output variance in a strongly time-dependent manner, with distinct parameters controlling transient and steady-state behavior for potassium and sodium conductances. At the level of neuronal excitability, sampling hundreds of thousands of parameter sets produced a heterogeneous population of firing behaviors, including non-firing, phasic, regular, and spontaneous activity. Across stimulus amplitudes, the dominant firing mode was a single spike at stimulus onset. At the same time, the regularly firing subpopulation exhibited a broad distribution of firing rates, with a mean that matched the classic Hodgkin-Huxley prediction. In the phasic subpopulation, action potential propagation and conduction velocity varied widely yet remained consistent with experimental ranges. Finally, global sensitivity analysis during spiking shows uniformly small first-order indices but large total-order indices, indicating that excitability is primarily governed by strong interactions among parameters rather than by any single conductance or kinetic parameter. These results support a population-based view of conductance-based modeling in which biologically relevant behavior emerges from structured regions of parameter space.

**Author summary:** Neurons are often modelled by first fitting ion-channel data to a few parameters, then integrating them into a single-neuron model. This typical method masks the fact that actual experiments show variability and that many different parameter sets can yield similar electrical behavior. In this research, we explored what happens when we keep rather than average out that variability. We revisited Hodgkin and Huxley’s classic squid giant axon studies and derived ranges for sodium and potassium channel parameters by resampling digitized points from the original Hodgkin-Huxley figures using bootstrap resampling. We then ran simulations of hundreds of thousands of squid axon models, each with a unique, experimentally grounded parameter set, and analyzed the collective results. This showed that the most common response was a single action potential rather than repetitive firing, aligning with the axon’s role in the rapid escape response. Additionally, we discovered that no single parameter alone controls spiking; rather, it depends on interactions among multiple parameters. Our findings advocate a practical change in biophysical modeling: instead of hunting for a single best-fit model, researchers should estimate parameter uncertainty directly from data, create large ensembles that sample this uncertainty, and perform sensitivity analyses on these ensembles before choosing any model for further study.

## Introduction

One of the primary approaches to unraveling the computational properties of neurons is the development of mathematical and computational models [1–7]. These models range from highly abstract representations that treat neurons as point-like integrators to biologically detailed simulations that incorporate the full complexity of a neuron’s structure and biophysics [7–10]. The latter, biologically detailed models, have become increasingly feasible due to advances in experimental techniques, computational resources, and optimization techniques [11–14]. These models aim to recreate the electrical and chemical environment of neurons, enabling researchers to simulate how neurons process information under various conditions. Our research has focused on building detailed models of Layer 5 pyramidal neurons [15–18] and voltage-gated ion channels [19–23] using multiple computational tools, while also critiquing the current state of this vital field [24].

The origins of biologically detailed modeling can be traced back to the seminal work of Hodgkin and Huxley, whose formulation of action potential generation in the squid axon laid the groundwork for the field [25]. Their experimental and theoretical framework established a systematic path from voltage-clamp recordings to differential equations describing ionic currents, culminating in spiking neuron models. Despite its enduring influence, this formalism functions as a one-way pipeline. Each step reduces complexity by fitting data to a small set of parameters while discarding the biological variability that is an inherent part of neuronal excitability.

The consequences of this reduction are profound. Hodgkin–Huxley–based models yield a single deterministic representation of a neuron, without accounting for biological variability. In our view, the most significant loss happens during the extraction of channel kinetics from voltage-clamp recordings. The kinetic rate constants are derived through curve fitting of data collected from multiple voltage-clamp recordings. Essentially, curve fitting aggregates variability into point estimates, collapsing biologically meaningful variance. Therefore, the resulting rate constants reflect the average of the voltage-clamp data. Combining several such deterministic models (for sodium, potassium, and other conductances) creates a model for neuron firing. In practice, many distinct parameter sets can generate equally valid spiking behavior [26], yet the standard pipeline collapses this diversity into a single point estimate. This means that conventional models excel at finding the best single fit but offer limited guarantees of robustness. Thus, at the end of the Hodgkin-Huxley analysis pipeline, we obtain a particular model of neuronal firing. Still, we do not know whether it is robust, stable, or representative. It has been demonstrated that by sampling the entire range of parameter variance in the Hodgkin-Huxley model, a family of spiking models can be generated [26]. In addition, many studies have demonstrated that variability in model parameters should not be overlooked when simulating neuronal behavior [12,13,27–47].

Moreover, biologically detailed models pose a multilayer problem [24,48,49]. They are constructed from neuronal morphologies, passive membrane parameters, and voltage- and ligand-gated ion channels. Some parameters crucial for describing ion channel kinetics may be irrelevant to a model of neuronal firing. Parameter degeneracy occurs because neuronal firing is simulated by numerically integrating a large set of differential equations. Thus, a set of equations providing a reasonable model of a voltage-gated ion channel may include parameters that are not important for simulating neuronal firing. Highly insightful work has been devoted to elucidating the variability and robustness of biologically detailed models [27,31,43–45,50–54]. However, variability analysis has been performed post-hoc, rather than as part of the model generation process [12,31,39,41,43,45,47,50]. We therefore asked a fundamental question: Is neuronal excitability in conductance-based models a property of single optimized parameter sets, or an emergent feature of interacting parameter populations shaped by biological variability? The considerable progress made over the past two decades, combined with the immense expansion in computer power, gives us hope that biologically detailed models may become a reality. This hope has driven our research in the past decades [15–18,20,23,24,55,56] and is at the core of the current work.

Here, we address this question by explicitly reintroducing parameter variability into the Hodgkin–Huxley modeling pipeline and embedding uncertainty and sensitivity analysis directly into model construction. We first extracted the variability of sodium and potassium channel kinetic parameters from the original Hodgkin–Huxley data using bootstrap resampling. We then propagated this experimentally grounded variability through a spatially extended Hodgkin–Huxley cable model of the squid giant axon using large-scale Monte Carlo simulations. By combining voltage-clamp–level analysis with simulations of action potential initiation and propagation, we quantified how parameter variability and interactions shape neuronal excitability across modeling levels. Finally, we applied global variance-based sensitivity analysis to identify how individual parameters and their interactions contribute to firing behavior, thereby linking channel kinetics, neuronal firing patterns, and robustness within a unified population-based framework. Together, these analyses show that neuronal excitability in the Hodgkin–Huxley framework is best understood as an emergent property of interacting parameter populations rather than as the outcome of a single optimized parameter set, and that biologically relevant firing behavior naturally arises from this variability.

## Methods

### Biophysical model of an excitable cable

Simulations of neuronal excitability, unless stated otherwise, used a spatially extended Hodgkin–Huxley–type cable model. The axon was represented as a one–dimensional cylindrical cable of length 10 cm and radius 0.025 cm, electrically discretized into 80 isopotential compartments. Each compartment contained voltage–gated sodium and potassium conductances as well as a passive leak conductance. The membrane potential *V(x,t)* evolved according to the cable equation with active ionic currents. The membrane dynamics were governed by:

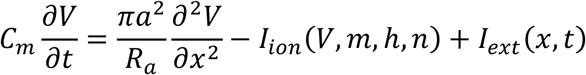

Here *C_m_* is the membrane capacitance per unit area (ranges defined in Table 1), *a* is the cable radius (0.025 cm in all simulations) , *R_a_* is the axial resistivity (35.4 Ω·cm in all simulations), *I_ion_* is the sum of ionic currents, and *I_ext_* represents externally injected current. The total ionic current density was the sum of fast sodium, delayed–rectifier potassium, and leak currents:

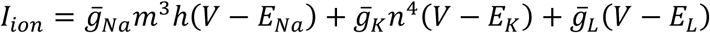

**Table 1:** Parameters of the Hodgkin-Huxley model and the ranges used in simulations. . The original value from the Hodgkin-Huxley manuscript is provided for comparison. The kinetic parameters obtained from the Bootstrap analysis are reported with the standard deviation from that analysis (and also in Eqs 1-6). Maximal conductance, reversal potentials, and membrane capacitance were extracted from Table 3 in the Hodgkin-Huxley manuscript [25].

| Name | Units | Original Value | Fitted Value | Standard Deviation |
| --- | --- | --- | --- | --- |
| $\bar{g}_K$ | mS/cm<sup>2</sup> | 36 | - | 6 |
| $E_K$ | mV | -77 | - | 2 |
| $A_{\alpha,n}$ | 1/(mV•ms) | 0.01 | 0.0092 | 0.0004 |
| $V_{1/2\alpha,n}$ | mV | 55 | 59.0 | 2.8 |
| $z_{\alpha,n}$ | mV | 10 | 6.9 | 1.8 |
| $A_{\beta,n}$ | 1/ms | 0.055 | 0.066 | 0.003 |
| $z_{\beta,n}$ | mV | 80 | 100.0 | 13.5 |
| $\bar{g}_{Na}$ | mS/cm<sup>2</sup> | 160 | - | 50 |
| $E_{Na}$ | mV | 45 | - | 6 |
| $A_{\alpha,m}$ | 1/(mV•ms) | 0.1 | 0.103 | 0.008 |
| $V_{1/2\alpha,m}$ | mV | 40 | 39.1 | 4.1 |
| $z_{\alpha,m}$ | mV | 10 | 6 | 2 |
| $A_{\beta,m}$ | 1/ms | 0.11 | 0.12 | 0.08 |
| $z_{\beta,m}$ | mV | 18 | 15.9 | 4.2 |
| $A_{\alpha,h}$ | 1/ms | 0.0027 | 0.0044 | 0.0014 |
| $z_{\alpha,h}$ | mV | 20 | 22.1 | 2.1 |
| $A_{\beta,h}$ | 1/ms | 1 | 0.96 | 0.10 |
| $V_{1/2\beta,h}$ | mV | 35 | 34.8 | 5.7 |
| $z_{\beta,h}$ | mV | 10 | 11.8 | 4.7 |
| $C_m$ | μF/cm<sup>2</sup> | 0.91 | - | 0.2 |
| $\bar{g}_L$ | mS/cm<sup>2</sup> | 0.3 | - | 0.1 |
| $E_L$ | mV | -53 | - | 4 |

Where 𝑔̅_𝑁𝑎_, 𝑔̅_𝐾_, and 𝑔̅_𝐿_ are maximal conductances and 𝐸_𝑁𝑎_, 𝐸_𝐾_, and 𝐸_𝐿_ are reversal potentials. Gating variables *m, h*, and *n* followed first–order voltage–dependent kinetics:

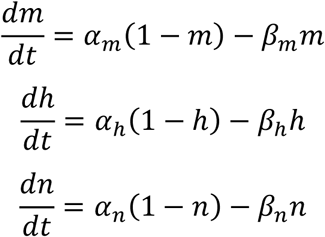

The transition rates were defined using standard Hodgkin–Huxley functional forms, with parameters controlling amplitudes, slopes, and voltage offsets (see Results below). All rate expressions were evaluated with numerical safeguards to avoid singularities, and gating variables were constrained to the interval [0,1].

### Spatial discretization and boundary conditions

The cable was discretized using a second–order finite–difference approximation of the axial second derivative. Sealed–end boundary conditions were imposed at both ends of the cable, enforcing zero axial current flow. This resulted in a system of 320 coupled ordinary differential equations representing membrane voltage and gating dynamics across space. An external current was injected into a single compartment located 0.4 cm from the cable’s sealed end. The injected current was specified as a total current and converted into a current density by dividing by the membrane surface area of the stimulated segment. The stimulus consisted of a square pulse with linear onset and offset ramps, beginning at 10 ms and ending at 80 ms. Voltage responses were recorded both at the injection site and at a distal location 8 cm away.

For each parameter set, the initial condition is chosen as the equilibrium (resting) state of the single–compartment Hodgkin-Huxley model corresponding to those parameters. The resting potential V_rest_ was defined as the voltage at which the total ionic current vanished when all gating variables were at steady state. In practice, we numerically found the solution to the algebraic function:

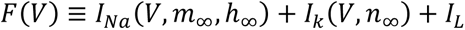

The steady-state gating variables were defined as: x_∞_(V)=α_x_(V)/(α_x_(V)+β_x_(V)) for x ∈ {m, h, n}. The equilibrium voltage was obtained using a damped Newton iteration with a finite-difference derivative approximation. This approach modified the standard method in three key ways. Step Damping: Instead of taking the full Newton step, only 30% of the step was applied. This smaller, more conservative update prevents the solver from jumping too far in any single iteration. Step Limiting: The maximum change in voltage per iteration was capped at 5 mV. This prevents large, unstable jumps when the local derivative is very small. Boundary Constraints: The voltage was strictly bounded within a physiological range (−100 mV to +50 mV) at each step. These modifications ensure that the algorithm remains stable and reliably converges to the correct physiological resting potential, even when starting from a broad range of initial guesses or exploring a wide parameter space in Monte Carlo simulations. Once V_rest_ was found, all compartments were initialized to this voltage and the corresponding steady-state gating values, yielding a spatially uniform resting state.

### Numerical integration and implementation

All simulations were implemented in Python (3.12.4) using NumPy (2.2.1) and JAX (0.8.1 [57]) for numerical computation, and the Diffrax library (0.7.0 [58]) for ordinary differential equation integration. The coupled system was integrated using an adaptive fifth–order Runge–Kutta method with relative and absolute tolerances of 10⁻⁵ and 10⁻⁶, respectively.

Simulations were run for 100 ms of biological time, and membrane potentials were recorded at fixed intervals of 0.1 ms. To enable large–scale parameter exploration, simulations were vectorized and parallelized using JAX’s just–in–time compilation, automatic vectorization, and parallel mapping across CPU cores.

Model parameters were sampled using simple Monte Carlo sampling over the ranges obtained in Figure 1. The sampled parameters included maximal conductances, reversal potentials, membrane capacitance, and all kinetic parameters governing sodium and potassium channel gating. The number of parameter sets sampled is indicated in the results and figure legends. Several safeguards were implemented to ensure numerical stability and to prevent unphysical states during large-scale Monte Carlo simulations. Rate expressions that contain potentially ill-conditioned terms of the form in the denominator were evaluated using limiting expressions when the denominator approached zero. All gating variables were constrained by clipping to the interval [0,1] at every solver evaluation.

**Figure 1:**
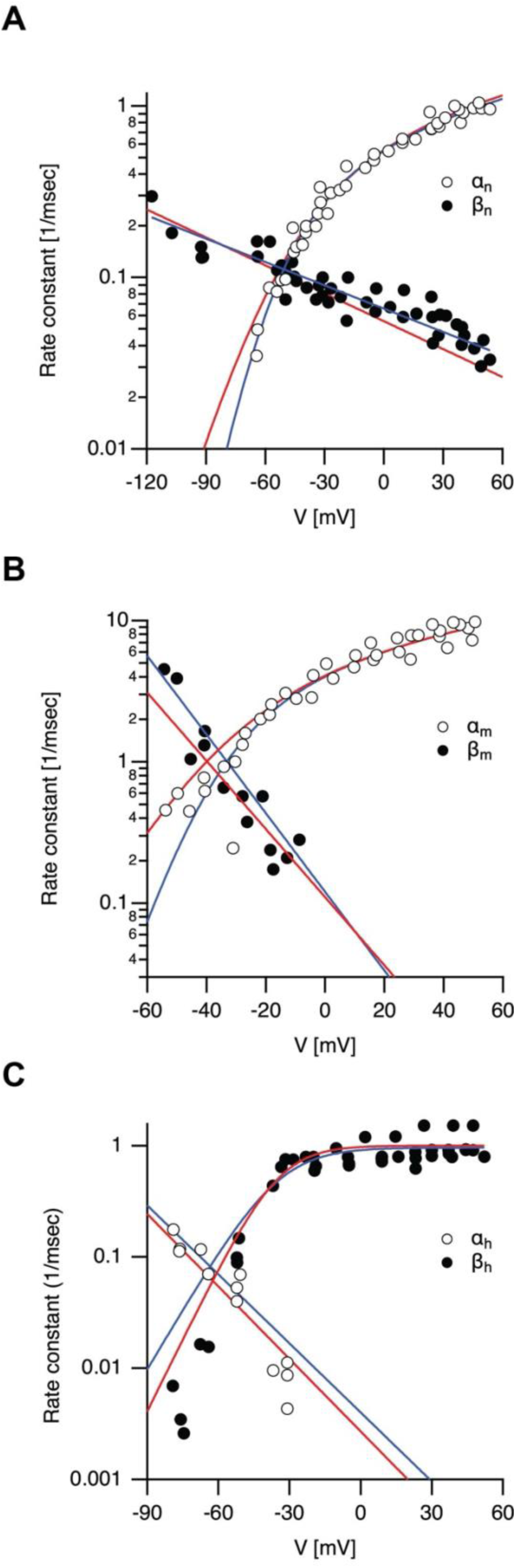
Refitting the rate constants of the Hodgkin-Huxley model. A, the forward (white circles) and backward (black circles) rate constants for the potassium conductance, extracted from the Hodgkin-Huxley paper. The original Hodgkin-Huxley-calculated rate constants are plotted in red, and the new fit from the Bootstrap analysis is plotted in blue. B, the same for the rate constants defining the activation of the sodium conductance. C, the same for the rate constants defining the inactivation of the sodium conductance.

As detailed above, the resting potential was computed using a damped Newton iteration to solve the steady-state current-balance equation. Newton updates were bounded in magnitude to avoid instability in poorly conditioned parameter regimes, and the finite-difference estimate of the derivative was regularized by enforcing a small minimum magnitude. Although the production simulations used an adaptive solver, this bound provided a robust initial step size for the stiffest parameter combinations encountered in the Monte Carlo ensemble.

### Sensitivity analysis

Sensitivity analysis evaluates how uncertain parameters influence the variability of model outputs. Numerous sensitivity measures are available [59–61]. In this study, we employed a global variance-based sensitivity analysis that calculates Sobol sensitivity indices, a well-established approach [62]. This global and non-intrusive method allows us to investigate interactions among model parameters [61]. To quantify how variability in Hodgkin–Huxley model parameters influences neuronal excitability, we performed a global variance-based sensitivity analysis. This approach explicitly treats model parameters as random variables and evaluates how their uncertainty propagates to variability in model outputs. Sensitivity analysis was embedded directly into the simulation pipeline rather than applied post hoc to a single optimized parameter set.

Each sampled parameter set was used to simulate the neuronal response to a brief suprathreshold current injection designed to elicit, when possible, a single action potential. From each simulation, scalar summary features were extracted to characterize excitability, including the peak membrane potential, action potential amplitude relative to rest, and time to peak depolarization. Parameter sets that failed to generate an action potential were retained in the analysis and assigned subthreshold output values.

Formally, a model output 𝑌 = 𝑓(𝑄_1_, 𝑄_2_, … , 𝑄_𝑝_) was defined as a function of *p* uncertain parameters *Q_i_*, including maximal conductances, reversal potentials, membrane capacitance, and kinetic parameters governing sodium and potassium channel gating. Sensitivity was quantified using Sobol variance decomposition.

The first-order Sobol sensitivity index for parameter *Q_i_* was defined as:

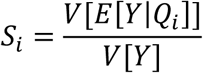

𝐸[𝑌|𝑄_𝑖_] represents the expected value of the output Y when the parameter Q_i_ is held constant. 𝑉[𝐸[𝑌|𝑄_𝑖_]] quantifies the variance explained by Q_i_ alone, and V[Y] is the variance of the output. The first-order Sobol sensitivity index indicates how much the model’s variance is expected to decrease when parameter Q_i_ is held constant. For example, if S_i_ =0.3, then 30% of the output variance is explained by Q_i_ alone. The sum of all first-order Sobol sensitivity indices cannot exceed one and equals one only if there are no interactions between parameters [63].

To capture interaction effects between parameters, total-order sensitivity indices were also computed. The total sensitivity index for parameter *Q_i_* was defined as:

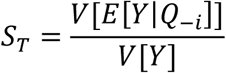

𝐸[𝑌|𝑄_−𝑖_] represents the expected value of the output Y when all parameters, except Q_i,_ are held constant. 𝑉[𝐸[𝑌|𝑄_−𝑖_]] quantifies the variance explained by all variables except Q_i_. For example, if S_Ti_ =0.5, then *Q_i_* (including its interactions with other variables) explains 50% of the output variance.

Sobol indices were estimated using Monte Carlo sampling based on Saltelli’s extension of the Sobol method [60]. Parameter values were sampled independently from distributions spanning ranges derived from Bootstrap analysis. All simulations used identical solver tolerances, temporal discretization, and spike-detection criteria to ensure numerical consistency across the ensemble. Sensitivity indices were computed separately for each output feature, allowing us to distinguish parameters that primarily influence spike initiation, spike amplitude, and spike timing. All simulations were carried out using custom Python code and the SALib sensitivity analysis library (1.5.1 [64]). The code is available in a GitHub repository at the address https://github.com/alon67/HodgkinHuxleyVariability.

## Results

The Hodgkin-Huxley analysis process has two main stages. First, a kinetic model for each channel is chosen and fitted to voltage-clamp currents, yielding numerical values for the rate constants. In the second step, these rate constants are plotted against membrane potential and fitted to a phenomenological equation that describes their voltage dependence. To study variability in the Hodgkin-Huxley model, we reanalyzed data from the Hodgkin-Huxley papers to determine the range of parameter variations. Because we lack direct access to the raw voltage-clamp traces, we could only extract parameters that define the Hodgkin-Huxley model’s rate constants. These rate constants are based on simplified assumptions made during data analysis to describe the gating of potassium channels with four cooperative activation gates and sodium channels with three activation gates plus one inactivation gate. Therefore, we extracted the rate constants by examining Hodgkin and Huxley’s original figures. We extracted the figures from the PDF of the Hodgkin-Huxley manuscript [25], enlarged them on a computer monitor, calibrated the figure axes to the screen coordinates in Adobe Photoshop, and then measured each data point by hand. To verify this extraction method, we compared the obtained values to the sample data provided in the Hodgkin-Huxley paper in tables 1 and 2 [25]. This was done for the six rate constants defining the Hodgkin-Huxley model. The voltage axis in the original Hodgkin-Huxley paper was relative to the resting membrane potential. We converted this to absolute membrane potential by adding a resting potential of −65 mV to the data and to the Hodgkin-Huxley equations. The original Hodgkin-Huxley equations describing the voltage dependence of the rate constants for potassium conductance are therefore:

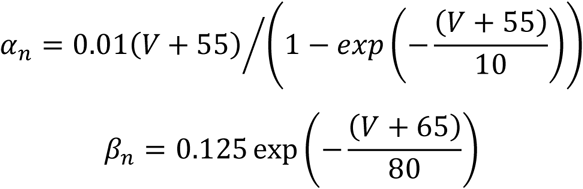

It is important to note that there is parameter degeneracy in the equation describing 𝛽_𝑛_. It can lead to multiple equivalent solutions during model investigation. Thus, it is better to simplify this rate constant to:

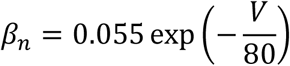

To estimate the parameter uncertainties in Equations 1 and 3, we digitized the potassium conductance rate-constant data from Figure 4 of the original Hodgkin–Huxley paper. Parameter uncertainty was quantified using a non-parametric bootstrap procedure implemented in Python. Specifically, the extracted data points were resampled with replacement to generate bootstrap datasets, which were refitted using the same nonlinear least-squares procedure as in the original fit. This yielded an empirical distribution for each fitted parameter, from which the mean value and standard deviation were computed and used as estimates of the parameter value and its variance. Parameter distributions for all rate constants analyzed in this work are shown in Supplementary Figure 1. The new rate constants with the mean fit and standard deviation are presented in Eqs. 1 and 2, and the fit is shown in Figure 1A.

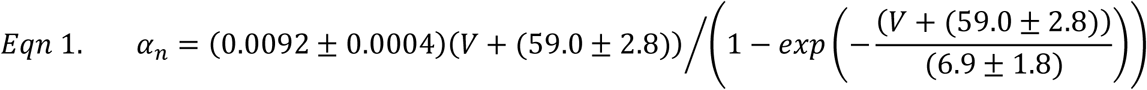

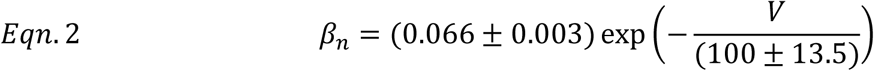

The sodium conductance has four rate constants, two for activation and two for inactivation. For the activation process, the rate constants from the Hodgkin-Huxley model, corrected for a resting membrane potential of −65 mV are:

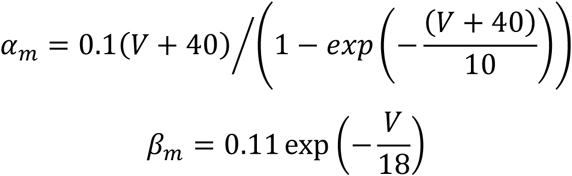

Extracting the activation rate constants from Figure 7 in the Hodgkin-Huxley paper and refitting them using bootstrap to estimate parameter variation provided:

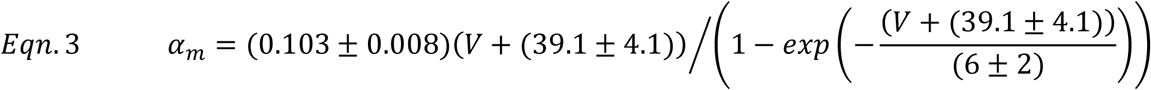

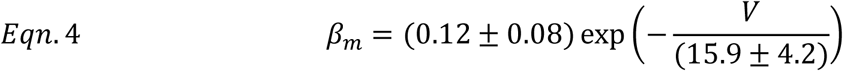

The new rate constants with the mean fit and standard deviation are presented in Eqs. 3 and 4, and the fit is shown in Figure 1B. For inactivation, the rate constants from the Hodgkin-Huxley paper are:

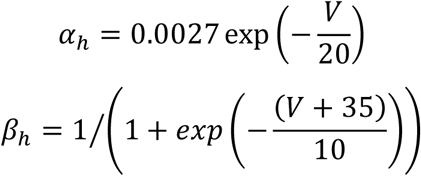

As with the previous rate constants, the data points were extracted from the original Hodgkin-Huxley paper (Figure 9). Bootstrap was used to estimate the standard deviation of the curve fit of these data to Eqs. 5 and 6, as shown in Figure 1C. All the parameters appearing in Eqs. 1-6 are summarized in Table 1, which also contains systematic naming of all the parameters.

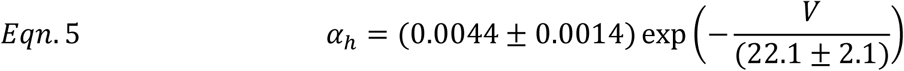

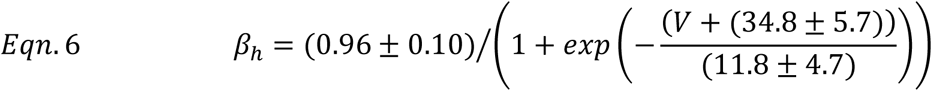

Almost all 15 parameters in this bootstrap analysis followed a bell-shaped distribution (Supp. Fig. 1). The two parameters controlling the backward rate of activation for the sodium conductance (𝛽_𝑚_) were an exception, likely due to the small number of data points and large scatter (Supp. Fig. 1I and 1J). Given this pattern, we decided to draw parameter values from Gaussian distributions for all Monte Carlo simulations. The upper and lower bounds of parameter sampling were set to the mean ± 2*std for each parameter, according to Eqs. 1-6 and Table 1. This rule generated negative values of 𝐴_𝛽,𝑚_ due to the asymmetric shape of its distribution (Supp. Fig. 1I). Therefore, the lower bound for this parameter was set to 10^-10^. The mean and standard deviation of the maximal conductances, reversal potentials, and membrane capacitance were obtained from Table 3 in the Hodgkin and Huxley paper [25]. Monte Carlo simulations in which parameter values were drawn from a uniform distribution yielded results similar to those obtained with parameter values drawn from a Gaussian distribution.

We used the results of this uncertainty analysis to assess parameter sensitivity in models of potassium and sodium conductances. We simulated each conductance under voltage-clamp conditions in a single compartment without spatial dimensions, and we computed the first-order Sobol’ sensitivity indices (Si) over time using Monte Carlo simulations. Figure 2A illustrates a voltage-clamp experiment where the membrane potential was stepped from −80 mV to −10 mV. The kinetic parameters defining equations 1 and 2 were randomly sampled as defined above. To focus solely on kinetic sensitivity, we simulated changes in the channel’s open probability without altering its maximal conductance. The potassium conductance, defined by five kinetic parameters in the Hodgkin-Huxley model, each influenced the output variance uniquely over time (Figure 2B).

**Figure 2:**
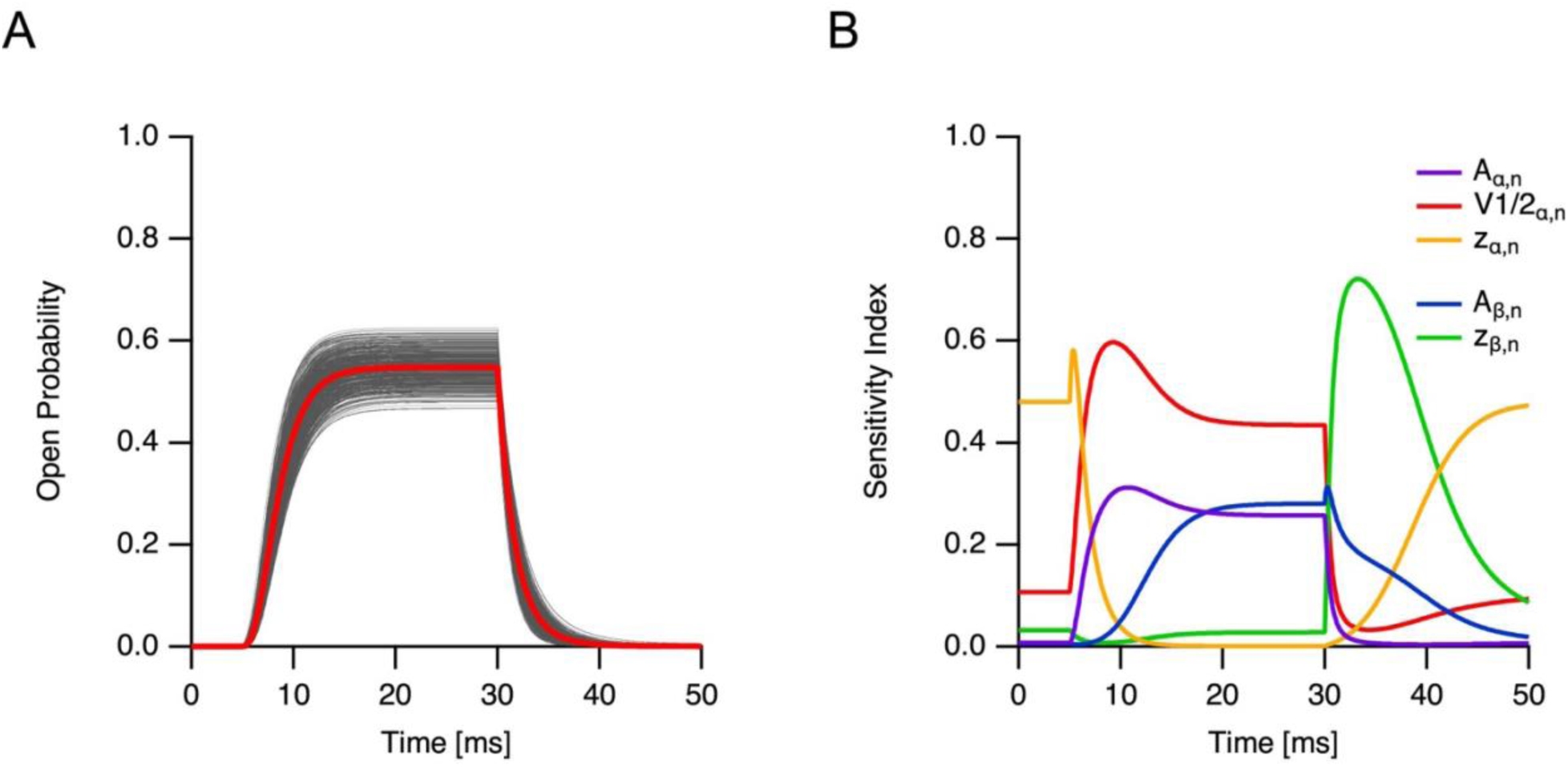
Sensitivity analysis of the potassium conductance. A, Simulations using different parameter sets of a voltage-clamp experiment in which the membrane potential was stepped from −80 to −10 mV. 500 simulations of the conductance open probability are shown in gray as a function of time, with the population average in red. B, First-order sensitivity indices, calculated from 98,304 Monte Carlo simulations (N=8192 with Saltelli sampling), for the five kinetic parameters defining the voltage dependence of the potassium conductance are plotted as a function of time.

Similarly, we performed Monte Carlo simulations to assess how the parameters defining the sodium conductance, as described by equations 3-6 and Table 1, affect its opening probability (Figure 3). Figure 3A shows a simulated voltage-clamp experiment in which the membrane potential was stepped from −80 mV to −10 mV. Randomly simulated traces are plotted, and the population average is overlaid in red. Ten kinetic variables define the sodium conductance in the Hodgkin-Huxley model. All of them displayed time-dependent sensitivity indices (Fig. 3B and 3C). Most parameters related to conductance activation showed transient changes when the voltage was stepped from −80 to −10 mV (Fig. 3B). As expected from the tenfold-slower inactivation process, the parameters controlling it exhibited slower time-dependent sensitivity indices.

**Figure 3:**
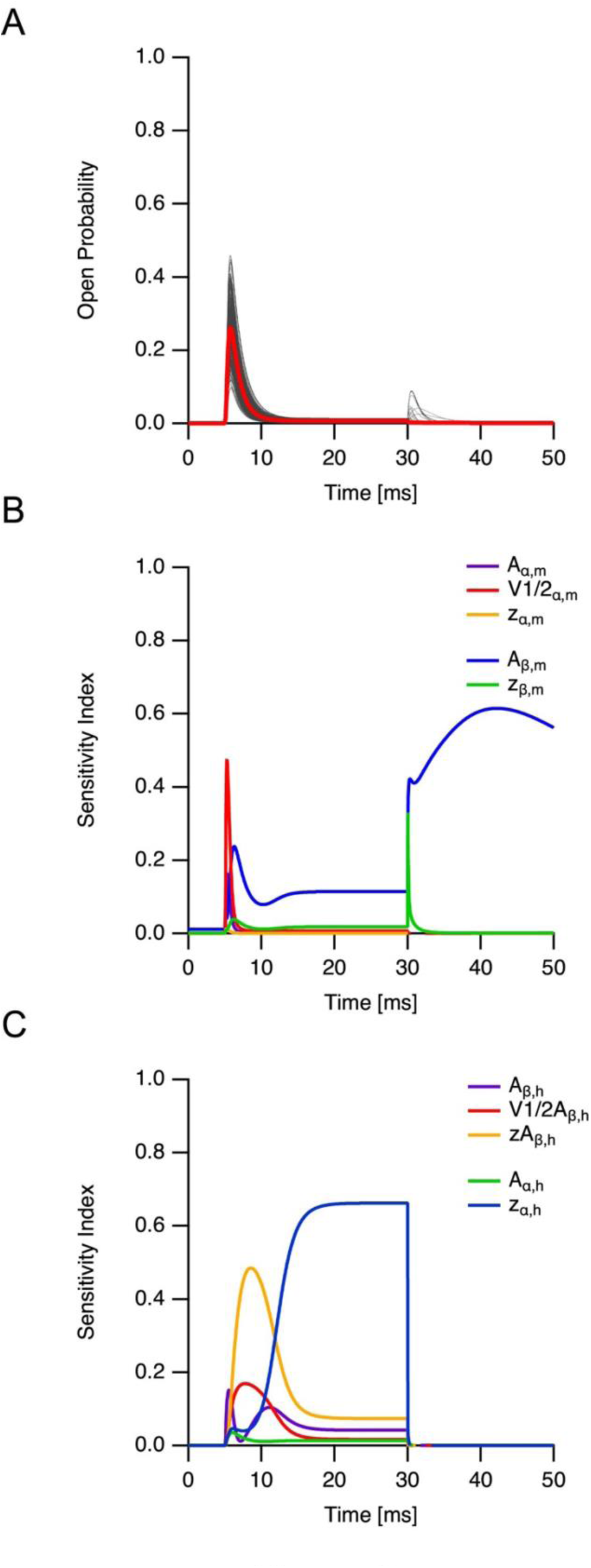
Sensitivity analysis of the sodium conductance. A, Simulations using different parameter sets of a voltage-clamp experiment in which the membrane potential was stepped from −80 to −10 mV. 500 simulations of the conductance open probability are shown in gray as a function of time, with the population average in red. B, First-order sensitivity indices, calculated from 90,112 Monte Carlo simulations (N=4096 with Saltelli sampling) , for the five kinetic parameters defining the voltage dependence of the sodium conductance activation process are plotted as a function of time. C, First-order sensitivity indices for the five kinetic parameters defining the voltage dependence of the sodium conductance inactivation process are plotted as a function of time.

The sensitivity analysis using Sobol’ first-order indices (Figures 2 & 3) revealed that all kinetic parameters affecting sodium and potassium conductances in the Hodgkin-Huxley model impact output variance. This was expected because the Hodgkin-Huxley model functions as a concerted system in which all transitions result in channel opening. In contrast, Markov models that include closed-closed transitions tend to exert less influence on the model’s output sensitivity [56].

After analyzing parameter sensitivity under voltage-clamp conditions, we integrated sodium and potassium conductances with the Hodgkin-Huxley model’s structural parameters in a 10 cm cable with a 0.5 mm diameter. A 70-ms square current pulse was applied at 0.4 cm from the sealed end to minimize end effects. For most simulations, we randomly sampled 300,000 parameter sets from Gaussian distributions based on the ranges listed in Table 1. Additionally, to ensure comprehensive coverage of the parameter space, we conducted simulations with 800,000 and 1,200,000 parameter sets, yielding results identical to those obtained with 300,000 sets (Supplementary Figure 2).

Figure 4A shows 500 individual simulations from this population, each receiving an 5 µA current injection through the electrode. The population average across all 300,000 simulations is shown in red and overlaid on these traces. Based on their firing patterns, these traces were categorized into four subpopulations: traces in which no APs were generated at the injection site throughout the entire 100 ms simulation duration (Fig. 4B), traces characterized by exactly one action potential occurring after stimulus onset (Fig. 4C), traces showing two or more action potentials during the stimulus period (Fig. 4D), and traces exhibiting one or more spontaneous action potentials during the initial 10 ms period before stimulus onset (Fig. 4E). Action potentials were identified using an upward threshold-crossing method with a detection threshold of −20 mV. This automatic method for selecting action potentials was verified by manually inspecting 500 random traces from each subcategory. Overall, 7.2% of the simulations exhibited spontaneous firing (Fig. 5). When no current was injected, the remaining population did not fire action potentials (Fig. 5, top left corner). As the current increased, the proportion of phasic and regular firing subpopulations grew (Fig. 5). The subthreshold response nearly disappeared as the current reached 8 µA. Interestingly, the regular-firing subpopulation increased to ∼14%, whereas most simulations (77.7% at 8 µA) produced a phasic response. We then examined the axon subpopulation that fired repeatedly during stimulation. Figure 6A shows three firing rate histograms of this subpopulation at different current levels. Within the parameter ranges in our bootstrap analysis, each current injection produced a wide distribution of regular firing frequencies. Their mean and standard deviation are plotted in Figure 6B as a function of the injected current. For comparison, we simulated the original Hodgkin-Huxley model parameters. We calculated the firing rate as a function of current. As expected from the similarity between the mean parameters we extracted in the Bootstrap analysis and the original Hodgkin-Huxley parameters (Table 1), these firing frequencies (red symbols in Fig. 6B) matched the mean firing rate of the regularly firing subpopulation.

**Figure 4.**
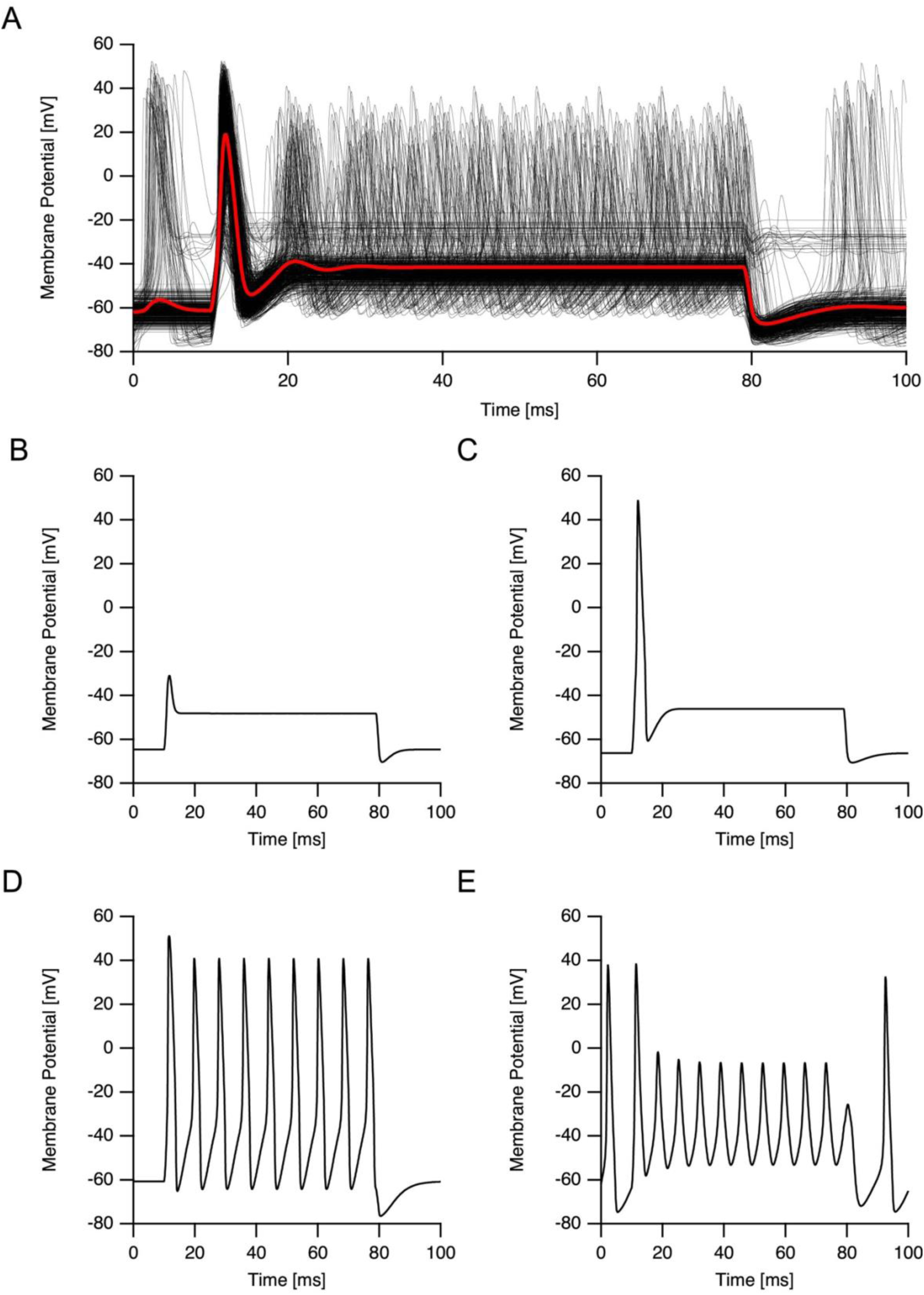
Population structure of firing behaviors in the Hodgkin-Huxley axon model. A, Five hundred representative membrane potential traces recorded at the site of current injection during a 70-ms square current pulse, randomly selected from the entire Monte Carlo ensemble of 300,000 parameter sets. Individual simulations are shown in gray, and the population-average membrane potential is overlaid in red. B–E, Based on their voltage responses, simulations were classified into four distinct firing phenotypes: B, nonfiring responses showing only subthreshold depolarization, C, phasic responses generating a single action potential at stimulus onset, D, regular firing responses producing multiple action potentials during the stimulus, and E, spontaneous firing responses in which action potentials occurred in the absence of injected current.

**Figure 5:**
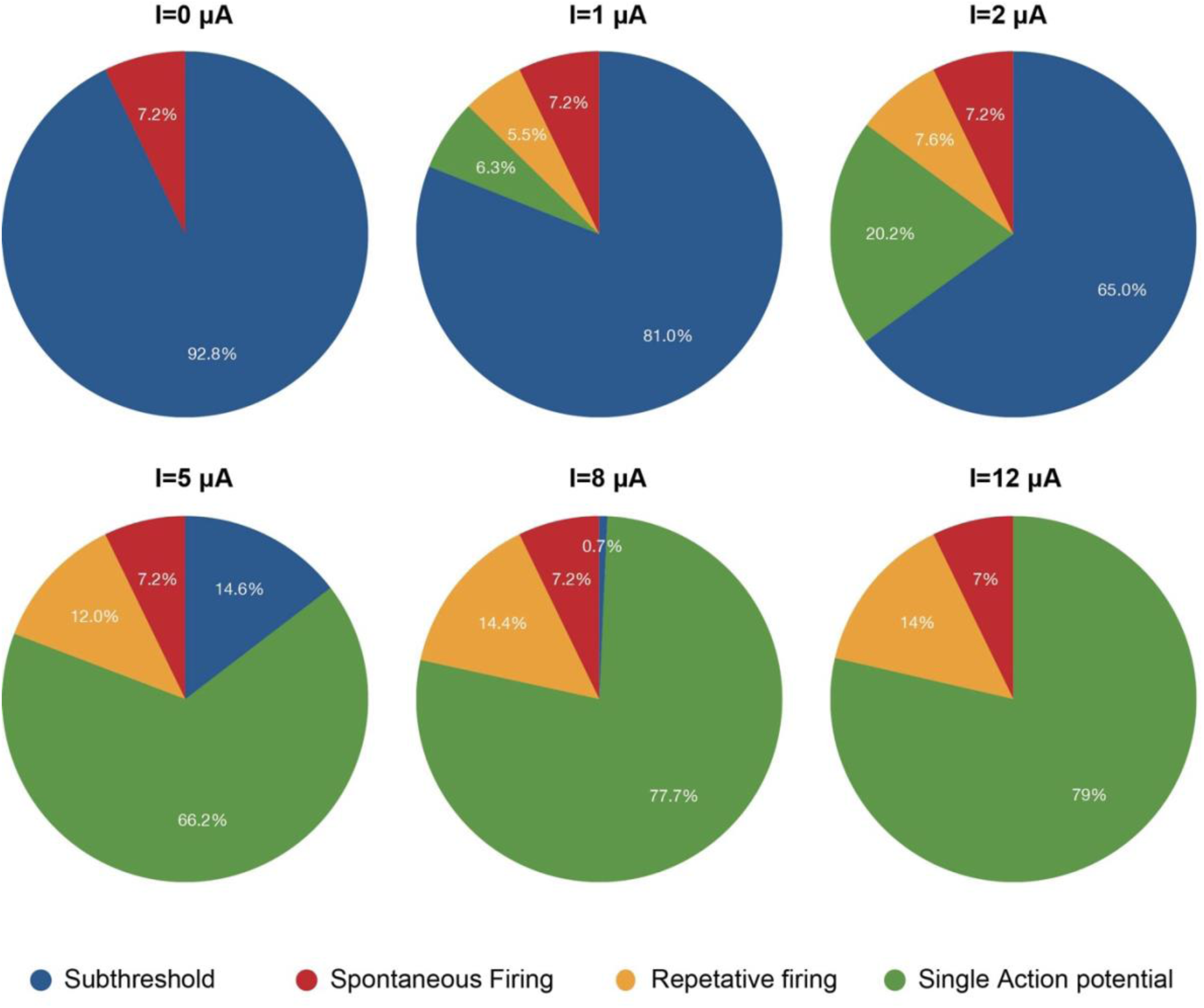
Population structure of firing behaviors in the Hodgkin-Huxley axon model as a function of injected current. Pie graphs show the relative numbers of each firing behavior observed in the Monte Carlo simulations. Subthreshold responses (blue), phasic responses that generate a single action potential at stimulus onset (green), regular firing responses that produce multiple action potentials during the stimulus (yellow), and spontaneous firing responses in which action potentials occur in the absence of injected current (red). The injected current is indicated above each pie graph.

**Figure 6.**
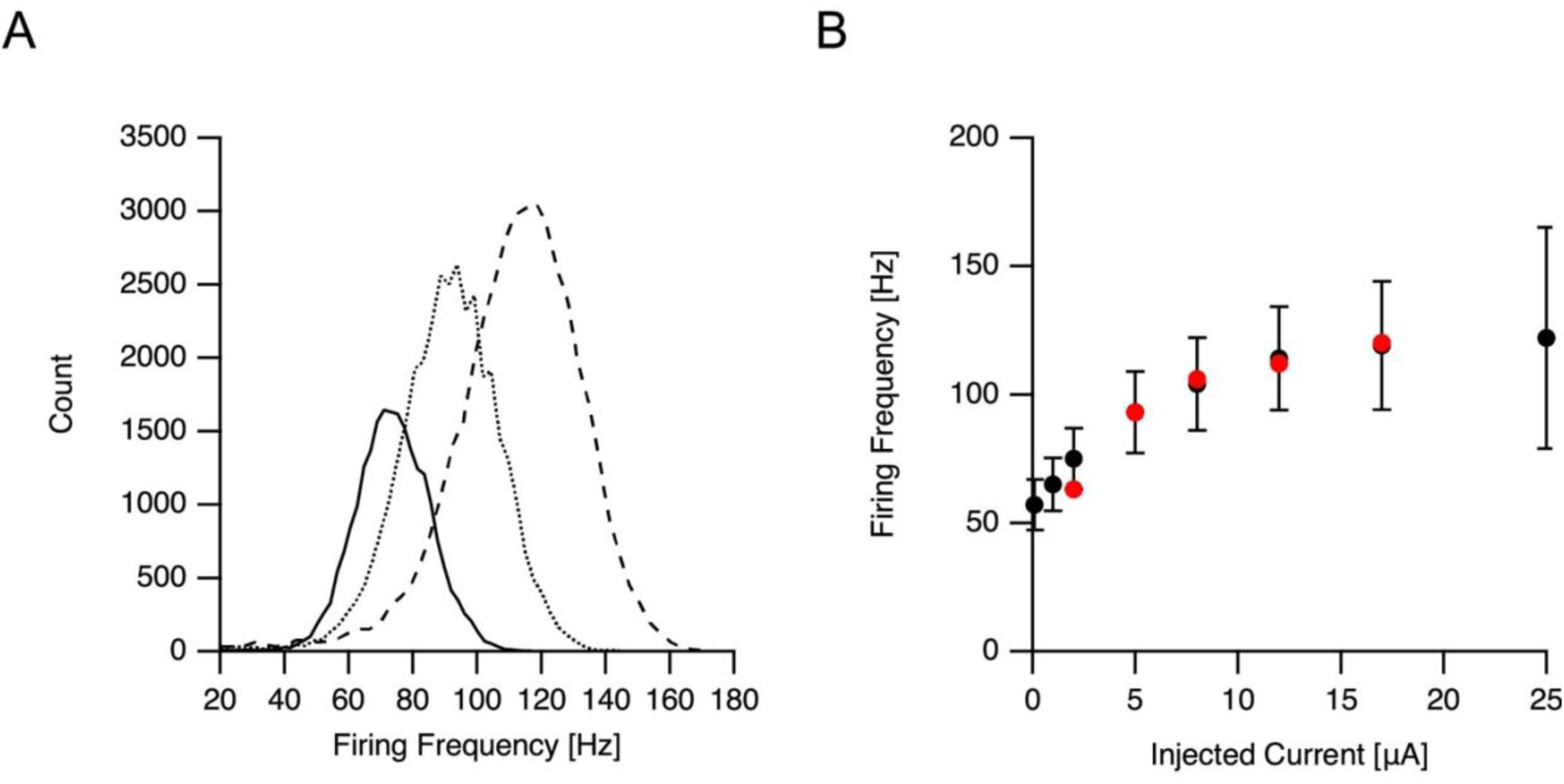
Firing rate variability in the regularly firing subpopulation. A, Histograms of firing rates for simulations classified as regularly firing, shown for three injected current amplitudes. Each histogram summarizes the distribution of firing frequencies across all parameter sets that produced sustained repetitive firing under the corresponding stimulus. The histograms were generated by current injections of 2 µA (lines), 5 µA (dots), and 12 µA (dashes). B, Mean firing rate (symbols) and standard deviation (error bars) of the regularly firing subpopulation plotted as a function of injected current amplitude. For comparison, the firing rates obtained from simulations using the original Hodgkin-Huxley parameter set are shown in red. The simulations using the original Hodgkin-Huxley parameter set were carried out in the same structural model and current injection levels as the rest of the simulations presented in this figure.

**Figure 7.**
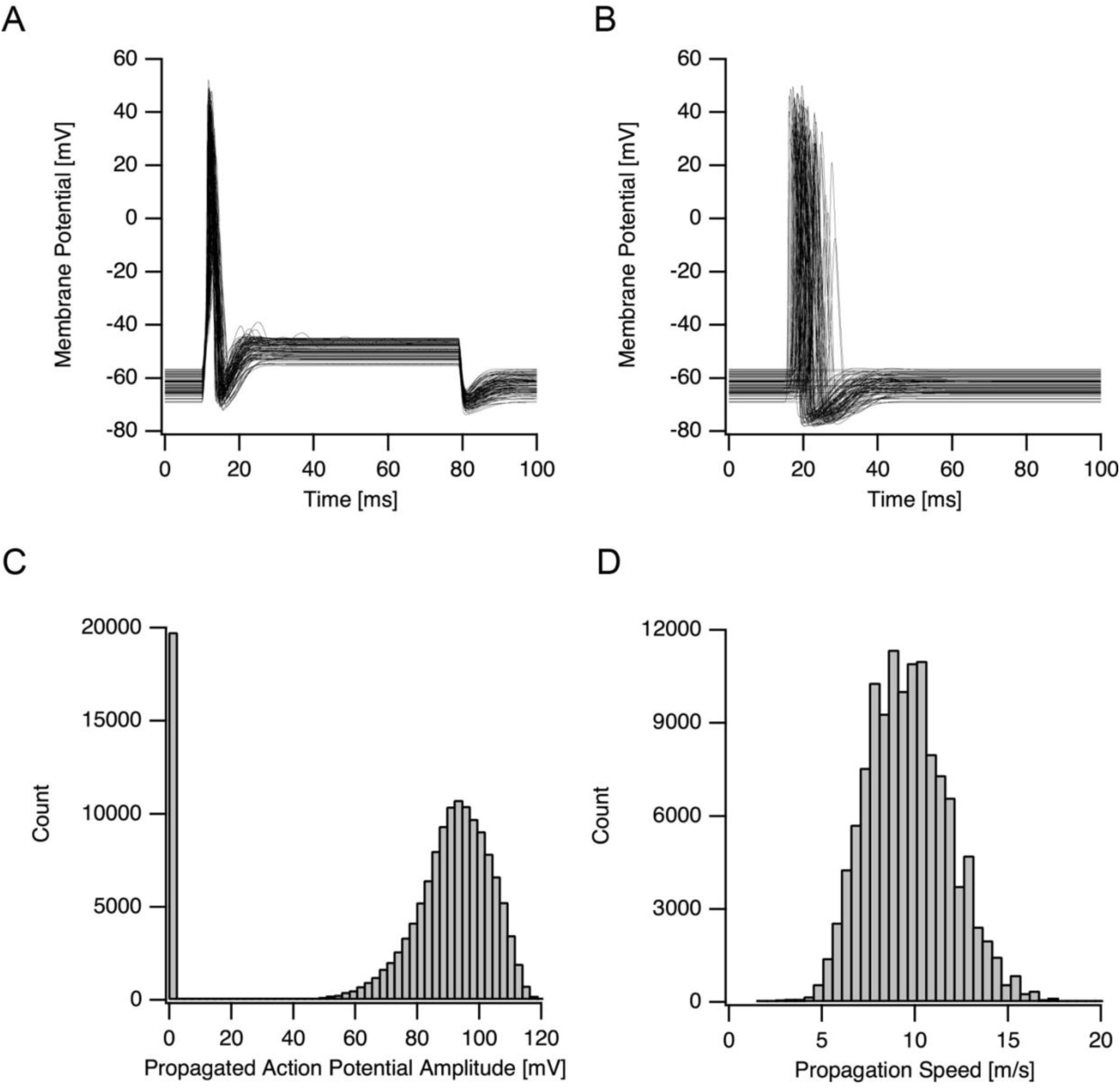
Propagation and variability of phasic action potentials in the axon model. A, Five hundred representative simulations from the phasic firing subpopulation, showing membrane potential traces recorded at the current injection site in response to a suprathreshold stimulus. B, Corresponding membrane potential traces recorded at a distal location 8 cm from the injection site, illustrating successful and failed propagation of action potentials along the cable. C, Distribution of peak action potential amplitudes at the distal recording site, reflecting variability in propagation efficacy, including complete propagation failures and attenuated spikes. D, Distribution of conduction velocities computed for successfully propagated action potentials.

The predominance of the phasic response in the simulated population was unexpected, given that the Hodgkin-Huxley model typically generates a regular firing pattern. We anticipated that the subpopulation characterized by regular firing would be the largest. However, as we will discuss later, the phasic response better reflects the physiological role of the squid’s giant axon [65–68]. Consequently, the subsequent analyses focused on simulations where phasic firing produced a single action potential.

We assessed whether the action potential propagated from its initiation site and measured its propagation speed. Figure 7A shows 500 overlaid simulations of phasic firing in response to a 5 µA injection at X = 0.4 cm. The propagated membrane potential at X = 8 cm is illustrated in Figure 7B. Successful propagation was defined by a distal action potential amplitude of 70 mV, while traces exhibiting subthreshold depolarization or decremental conduction below this threshold were classified as failed propagation. As expected, some action potentials successfully initiated at the current injection site but failed to reach the distal recording site, resulting in a broad distribution of amplitudes at the distal site (Fig. 7C). This data on action potential propagation can be used to refine parameter ranges. The conduction velocity was calculated by dividing the distance between the injection site and the distal recording site by the time delay between the action potential peaks at the two locations. The propagation speed in these simulations ranged from approximately 5 to 15 m/s (Fig. 7D), consistent with experimental data [67,69].

We calculated the first- and total-order sensitivity indices (Fig. 8) for the subpopulation of simulations exhibiting phasic firing (Fig. 7A), as these may be of physiological importance. These simulations were carried out in a point model of the neuron to reduce computational load. To validate these simulations, we performed the same analyses shown in Figures 4-6 on the point neuron firing patterns, yielding similar results. The simulation generated 376,832 traces (N=8192, Saltelli sampling for 22 parameters), out of which 182,386 displayed an action potential at stimulus onset. Sensitivity analysis presented in Figure 8 was performed on this population. As expected, the first-order sensitivity indices were small during both the action potential (at t= 12 ms, indicated by a blue arrow in Fig. 8A and 8B) and the subsequent sustained membrane potential (at t = 70 ms, indicated by a magenta arrow in Fig. 8A and 8B). The sustained potential was influenced mainly by two parameters of the potassium conductance (V1/2_α,n_ and A_β,n_), consistent with their role in the depolarization block following the initial firing of a single action potential (Fig. 7A). The low first-order indices highlight the minimal impact of individual parameters after numerical integration of the differential equations. In contrast, the total-order sensitivity indices were high for nearly all parameters during both the initial action potential and the sustained membrane potential (Fig. 8C and 8D), indicating strong interdependence among parameters due to their coupling with membrane potential.

**Figure 8.**
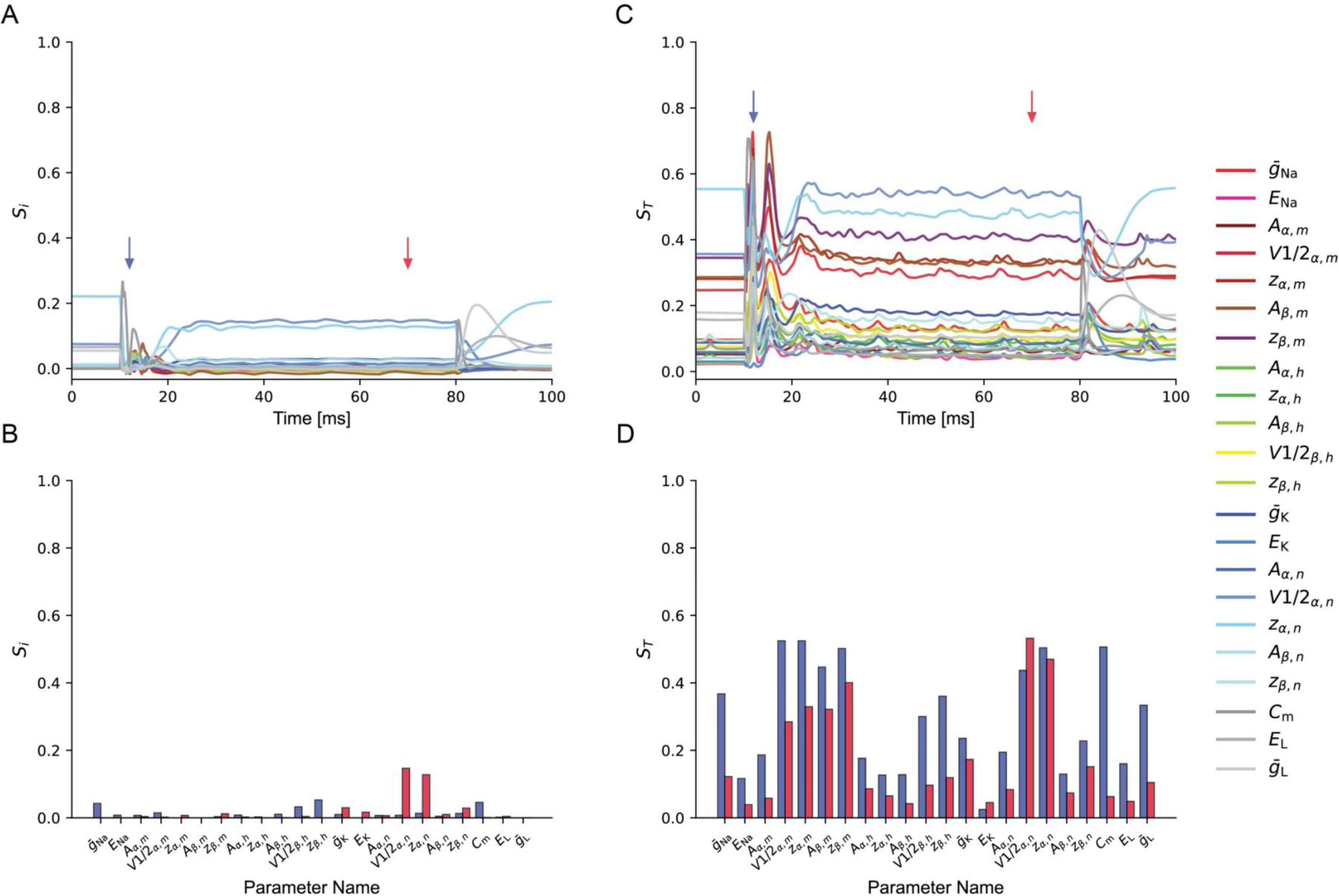
Global sensitivity structure of excitability in the phasic firing subpopulation. A, Time-dependent first-order Sobol sensitivity indices (S_i_) for selected model outputs during the action potential and the subsequent sustained membrane potential, computed across simulations classified as phasic firing. B, Values of S_i_ during the action potential at the injection site (indicated by a blue arrow in A) and during the sustained membrane potential later in the simulation (indicated by a magenta arrow in A). C, Time-dependent total-order Sobol sensitivity indices (S_T_) for selected model outputs during the action potential and the subsequent sustained membrane potential, computed across simulations classified as phasic firing. D, Values of S_T_ during the action potential at the injection site (indicated by a blue arrow in C) and during the sustained membrane potential later in the simulation (indicated by a magenta arrow in C). The simulation generated 376,832 traces (N=8192 in Saltelli sampling for 22 parameters), out of which 182,386 displayed an action potential at stimulus onset. Sensitivity analysis presented in the figure was performed on this population.

## Discussion

This study explores how explicitly including parameter variability in the Hodgkin–Huxley model alters our understanding of neuronal excitability and stability. We combined uncertainty analysis of channel kinetics from original data with large-scale Monte Carlo simulations and global sensitivity analysis integrated into the modeling process. Re-fitting potassium and sodium channel rates via bootstrap resampling revealed significant, uneven parameter uncertainty, indicating degeneracy in the voltage-dependent kinetics (Figure 1 and Table 1). Sensitivity analysis of voltage-clamp simulations showed that all kinetic parameters influence output variance over time, with different parameters dominating transient and steady-state behaviors of potassium and sodium channels (Figures 2 and 3). Incorporating these variable conductances into an excitable cable model and sampling hundreds of thousands of parameter sets produced diverse firing patterns: non-firing, phasic, regular, and spontaneous activity (Figures 4 and 5). The population displaying regular firing showed a broad range of firing rates, with the average aligning with the classic Hodgkin–Huxley result, emphasizing that typical behavior arises from a population, not a single solution (Figure 6). Further analysis of phasic responses showed considerable variability in action potential propagation and conduction velocity, consistent with experimental data from the squid giant axon (Figure 7). Global sensitivity analysis indicated that, while first-order sensitivity indices were small, total-order indices were large, suggesting that excitability depends mainly on interactions among parameters rather than on any single kinetic or conductance parameter (Figure 8).

In the present study, we applied uncertainty and global sensitivity analyses to the canonical Hodgkin–Huxley model of the squid giant axon to explicitly reintroduce biological variability into a framework that has traditionally produced a single deterministic solution. Our analysis was intentionally restricted to one well-defined instantiation of the Hodgkin–Huxley model, not as a claim of completeness, but as a controlled test case for examining how variability and parameter interactions shape excitability within this classical formalism. As emphasized in the Introduction, the Hodgkin–Huxley pipeline collapses experimental variability early in the modeling process. Here, we asked what emerges when that variability is preserved and propagated throughout the entire excitable system.

A first key result is that parameter sensitivity depends strongly on the level of description at which the model is interrogated. Under voltage-clamp conditions, individual kinetic parameters exert pronounced, time-dependent first-order effects on channel behavior (Figures 2 and 3). In this regime, sensitivity indices reflect the direct control of specific rate constants over activation and inactivation processes, closely mirroring the structure of the underlying equations. However, when these conductances are embedded in a spatially extended cable model and driven to spike, the situation changes qualitatively. During action potential generation, first-order Sobol indices were small (Figure 8A, 8B), indicating that no single parameter independently controls firing. At the same time, total-order indices remain large for nearly all parameters (Figure 8C, 8D), revealing that excitability is governed primarily by strong, high-order interactions mediated through membrane potential coupling. This contrast highlights a fundamental transformation in parameter influence when moving from isolated channel dynamics to integrated neuronal behavior.

These findings compel a reassessment of the “database” approach to biophysical modeling, which has traditionally focused on the variability of conductance densities while holding channel kinetics constant . The important work by Prinz et al. [44] demonstrated that similar network activity can arise from disparate sets of maximal conductances, establishing the concept of multiple solutions in neuronal modeling. While subsequent studies [41,50] expanded this framework to include morphological variability, voltage-dependent rate constants are typically treated as a fixed scaffold upon which conductance densities are tuned. This philosophy was paralleled in automated optimization, where genetic algorithms were used to constrain complex compartmental models. Studies by Vanier and Bower [70] and Keren et al. [17,55] showed that evolutionary algorithms could effectively find conductance densities that matched experimental firing patterns. Similarly, Druckmann et al. used multi-objective optimization to constrain conductance parameters for cortical interneurons [52] and pyramidal cells [51,71], treating the underlying channel kinetics as fixed building blocks. However, maximal conductances act primarily as linear scaling factors, whereas kinetic parameters determine the timing and non-linear state transitions of the system. Our global sensitivity analysis reveals that kinetic parameters, frozen in both database exploration and standard genetic optimization pipelines, are primary drivers of excitability through high-order interactions (Figure 8). This supports the observations of Goaillard et al. [47], who noted that ion channel degeneracy involves covariation in biophysical properties beyond simple conductance magnitudes. Consequently, modeling robustness solely through conductance variation ignores a fundamental dimension of biological variability, potentially underestimating the compensatory mechanisms available to excitable systems.

Ori et al. provide an elegant and highly instructive demonstration that, despite the apparent high dimensionality of the Hodgkin-Huxley model, the functional outcome of excitability can be organized in a much lower-dimensional physiological space [26]. Conceptually, this work anticipated the need to move beyond single parameter sets and to view excitability as a population-level property of the Hodgkin-Huxley model. In their analysis, they systematically rescaled the original Hodgkin-Huxley rate constants across a broad range of independent values. They showed that the resulting behaviors cluster along two composite axes that capture structural and kinetic contributions. They propose that slow sodium inactivation is a local homeostatic mechanism that stabilizes excitability within this reduced plane. In our manuscript, we approached the same overarching question from a complementary direction, by grounding parameter dispersion in a data-driven uncertainty analysis of the original voltage-clamp figures and using bootstrapping to estimate parameter-specific variability rather than imposing uniform rescaling across rate constants. While Ori et al. isolate a powerful geometric principle and mechanistic stabilizer using short-pulse stimulation in a single-compartment setting (7 µA, 1 ms), our results emphasize how experimentally constrained, parameter-specific variability and parameter interactions shape spike initiation, waveform, and propagation in an extended axon, and why kinetic parameters must be treated on equal footing with conductances when interpreting robustness and degeneracy in Hodgkin-Huxley-type models.

The population simulations further demonstrate that preserving parameter variability fundamentally alters the qualitative behavior of the model. Sampling hundreds of thousands of parameter sets produces a heterogeneous ensemble of responses, including non-firing, phasic firing, regular firing, and spontaneous activity (Figures 4 and 5). Notably, the dominant subpopulation consists of models that fire a single action potential in response to sustained depolarization (Figure 4C), while regularly firing models constitute a minority. At first glance, this result appears to contradict the behavior of the original Hodgkin–Huxley model, which typically produces repetitive firing under constant current injection. However, when interpreted in light of the squid giant axon’s physiological role, the predominance of phasic firing emerges as a biologically appropriate outcome rather than an anomaly.

The giant axon of the squid is a highly specialized structure that triggers rapid, all-or-none mantle contraction during escape behavior [65,66]. Experimental studies have shown that, under natural conditions, the axon typically fires a single action potential in response to sensory stimuli, particularly visual flashes in cold seawater, initiating a powerful jet-propelled response [65,66,68,72,73]. More complex patterns, one to three spikes or even complete silence, occur only in delayed escape behaviors involving chemical stimulation and coordination with the parallel small-axon system. This firing behavior reflects the axon’s intrinsic excitability, in which a neuron responds to sustained depolarization with a single spike followed by silence [74,75]. The predominance of phasic firing in our simulations, therefore, closely aligns with the known physiological function of the squid giant axon. This alignment suggests that the classic Hodgkin–Huxley parameter set represents a special case, rather than the axon’s physiological function, within a broader, biologically relevant parameter landscape.

Despite this convergence with physiology, the variability extracted in this study has clear limitations. Parameter uncertainty was estimated after selecting a specific kinetic model and was based on digitized data rather than original voltage-clamp recordings. As a result, the inferred variability is both incomplete, because it does not reflect alternative model structures, and likely inflated relative to true biological variability due to experimental scatter and fitting uncertainty. Nevertheless, even under these constraints, a large region of parameter space robustly predicts phasic firing and realistic propagation behavior, including conduction velocities within experimentally observed ranges (Figure 7D). This robustness suggests that meaningful functional predictions can emerge from population-based approaches even when uncertainty estimates are imperfect.

Importantly, not all parameters exhibit the same level of uncertainty. The bootstrap analysis reveals substantial heterogeneity in parameter variance (Table 1), reflecting differences in data density, voltage coverage, and scatter in the original Hodgkin–Huxley figures. While biological and experimental sources of variability cannot be disentangled here, this observation underscores the importance of explicit uncertainty analysis as a prerequisite for sensitivity analysis and model construction.

More broadly, the results presented here point toward an alternative paradigm for biophysical modeling. Rather than seeking a single optimized parameter set, our findings argue for a workflow in which uncertainty analysis is performed first, followed by generation of a large ensemble of models that is analyzed as a structured population. Within such populations, physiological behavior corresponds to regions of parameter space rather than isolated points. Selecting a single “best” model, whether by manual tuning or automated optimization, inevitably discards this structure and is somewhat analogous to overfitting. Even after constraining models to match physiological behavior, a nontrivial parameter manifold remains, and simulations must therefore account for variability rather than collapsing it into an average.

The parameters in the voltage-dependent rate equations tend to be correlated because the same data can often be fitted equally well by adjusting amplitudes, voltage offsets, slope factors, or balancing forward and backward rates. This practical degeneracy means many different parameter combinations can produce very similar activation and inactivation curves, as well as nearly identical channel kinetics. However, our findings indicate that these kinetic degrees of freedom remain crucial when analyzing neuronal excitability, especially at the level of spiking. While individual parameters generally have weak effects on their own after considering the full nonlinear system, their combined interactions have a significant impact. Therefore, even if correlations reduce the apparent dimensionality of the kinetic parameter space, they do not render these parameters irrelevant. Instead, they emphasize why sensitivity analysis must consider their coupled effects, as ignoring them would underestimate the importance of parameter interactions in determining neuronal excitability.

Finally, the present work also highlights directions for future extension. We did not include variability in axon diameter or morphology, factors known to influence excitability and conduction velocity strongly. Incorporating morphological uncertainty, additional conductances, or alternative channel formalisms such as Markov models will further enrich population-level analyses. Markov models, in particular, allow description of more complex kinetics than the Hodgkin–Huxley formalism, but also introduce transitions that may contribute weakly to functional sensitivity [56]. Applying the same uncertainty- and interaction-focused framework to such models will help clarify which aspects of added biophysical detail meaningfully shape neuronal behavior.

In summary, our results demonstrate that neuronal excitability in the Hodgkin–Huxley framework is best understood not as the property of a single parameter set, but as an emergent feature of a population defined by experimentally grounded variability and strong parameter interactions. Embracing this perspective is essential for constructing robust, physiologically interpretable biophysical models.

## Author Contribution

The author conceived the study, performed all simulations, analyzed the data, and wrote the manuscript. This manuscript is a one-man show.

## Data Availability

All simulations and analyses are publicly available on GitHub at https://github.com/alon67/HodgkinHuxleyVariability. No experimental data were generated for this study.

## Ethics Statement

No human or animal subjects were used in this study.

## References

1. Rall W. Electrophysiology of a dendritic neuron model. Biophys J. 1962;2: 145–167. 10.1016/s0006-3495(62)86953-7

2. Rall W. Distributions of potential in cylindrical coordinates and time constants for a membrane cylinder. Biophys J. 1969;9: 1509–41. doi:10.1016/S0006-3495(69)86468-4

3. Rall W. Theoretical Significance of Dendritic Trees for Neuronal Input-Output Relations. Reis RF, editor. Neural theory and modeling. Ojai: Stanford University Press, Palo Alto; 1964.

4. Segev I, Rall W. Excitable dendrites and spines: earlier theoretical insights elucidate recent direct observations. Trends Neurosci. 1998;21: 453–460. 10.1016/S0166-2236(98)01327-7

5. Hines ML, Markram H, Schürmann F. Fully implicit parallel simulation of single neurons. J Comput Neurosci. 2008;25: 439–448. doi:10.1007/s10827-008-0087-5

6. King JG, Hines M, Hill S, Goodman PH, Markram H, Schürmann F. A component-based extension framework for large-scale parallel simulations in NEURON. 2009;3: 1–11. doi:10.3389/neuro.11.010.2009

7. Markram H, Muller E, Ramaswamy S, Reimann MW, Abdellah M, Sanchez CA, et al. Reconstruction and Simulation of Neocortical Microcircuitry. Cell. 2015;163: 456–492. doi:10.1016/j.cell.2015.09.029

8. Segev I, Schneidman E. Axons as computing devices: basic insights gained from models. J Physiol Paris. 1999;93: 263–270. 10.1016/S0928-4257(00)80055-8

9. Hay E, Schürmann F, Markram H, Segev I. Preserving axosomatic spiking features despite diverse dendritic morphology. J Neurophysiol. 2013;109: 2972–81. doi:10.1152/jn.00048.2013

10. Carnevale NT, Hines ML. The NEURON book. Cambridge, UK ; New York: Cambridge University Press; 2006.

11. Birgiolas J, Haynes V, Gleeson P, Gerkin RC, Dietrich SW, Crook S. NeuroML-DB: Sharing and characterizing data-driven neuroscience models described in NeuroML. PLoS Comput Biol. 2023;19: e1010941. doi:10.1371/journal.pcbi.1010941

12. Schneider M, Bird AD, Gidon A, Triesch J, Jedlicka P, Cuntz H. Biological complexity facilitates tuning of the neuronal parameter space. PLoS Comput Biol. 2023;19: e1011212. doi:10.1371/journal.pcbi.1011212

13. Ladd A, Kim KG, Balewski J, Bouchard K, Ben-Shalom R. Scaling and Benchmarking an Evolutionary Algorithm for Constructing Biophysical Neuronal Models. Front Neuroinform. 2022;16. doi:10.3389/fninf.2022.882552

14. Deistler M, Kadhim KL, Pals M, Beck J, Huang Z, Gloeckler M, et al. Jaxley: Differentiable simulation enables large-scale training of detailed biophysical models of neural dynamics. 2024. doi:10.1101/2024.08.21.608979

15. Gurkiewicz M, Korngreen A. A numerical approach to ion channel modelling using whole-cell voltage-clamp recordings and a genetic algorithm. PLoS Comput Biol. 2007;3: e169. doi:10.1371/journal.pcbi.0030169

16. Almog M, Korngreen A. A quantitative description of dendritic conductances and its application to dendritic excitation in layer 5 pyramidal neurons. The Journal of Neuroscience. 2014;34: 182–96. doi:10.1523/JNEUROSCI.2896-13.2014

17. Keren N, Peled N, Korngreen A. Constraining compartmental models using multiple voltage recordings and genetic algorithms. J Neurophysiol. 2005/08/12. 2005;94: 3730–3742. doi:10.1152/jn.00408.2005

18. Ben-Shalom R, Liberman G, Korngreen A. Accelerating compartmental modeling on a graphical processing unit. Front Neuroinform. 2013;7: 1–8. doi:10.3389/fninf.2013.00004

19. Segev D, Korngreen A. Kinetics of two voltage-gated K+ conductances in substantia nigra dopaminergic neurons. Brain Res. 2007;1173: 27–35. doi:10.1016/j.brainres.2007.08.006

20. Almog M, Degani-Katzav N, Korngreen A. Kinetic and thermodynamic modeling of a voltage-gated sodium channel. European Biophysics Journal. 2022;51. doi:10.1007/s00249-022-01591-3

21. Almog M, Barkai T, Lampert A, Korngreen A. Voltage-Gated Sodium Channels in Neocortical Pyramidal Neurons Display Cole-Moore Activation Kinetics. Front Cell Neurosci. 2018;12: 187. doi:10.3389/fncel.2018.00187

22. Gurkiewicz M, Korngreen A, Waxman SG, Lampert A. Kinetic modeling of nav1.7 provides insight into erythromelalgia-associated F1449V mutation. J Neurophysiol. 2011;105: 1546–1557. doi:10.1152/jn.00703.2010

23. Ben-Shalom R, Aviv A, Razon B, Korngreen A. Optimizing ion channel models using a parallel genetic algorithm on graphical processors. Journal of neuroscience. 2012;206: 189–194. doi:10.1016/j.jneumeth.2012.02.024

24. Almog M, Korngreen A. Is realistic neuronal modeling realistic? J Neurophysiol. 2016;116. doi:10.1152/jn.00360.2016

25. Hodgkin AL, Huxley AF. A quantitative description of membrane current and its application to conduction and excitation in nerve. J Physiol. 1952;117: 500–544. 10.1113/jphysiol.1952.sp004764

26. Ori H, Marder E, Marom S. Cellular function given parametric variation in the hodgkin and huxley model of excitability. Proc Natl Acad Sci U S A. 2018;115. doi:10.1073/pnas.1808552115

27. Goldman MS, Golowasch J, Marder E, Abbott LF. Global structure, robustness, and modulation of neuronal models. J Neurosci. 2001;21: 5229–5238. doi:21/14/5229 [pii]

28. Hafner M, Koeppl H, Hasler M, Wagner A. ‘ Glocal ’ Robustness Analysis and Model Discrimination for Circadian Oscillators. 2009;5. doi:10.1371/journal.pcbi.1000534

29. Swensen AM, Bean BP. Robustness of burst firing in dissociated purkinje neurons with acute or long-term reductions in sodium conductance. J Neurosci. 2005;25: 3509–3520. 10.1523/JNEUROSCI.3929-04.2005

30. Weaver CM, Wearne SL. Neuronal firing sensitivity to morphologic and active membrane parameters. PLoS Comput Biol. 2008;4: 0130–0150. doi:10.1371/journal.pcbi.0040011

31. Achard P, De Schutter E. Complex parameter landscape for a complex neuron model. PLoS Comput Biol. 2006;2. doi:10.1371/journal.pcbi.0020094

32. Sobie E a. Parameter sensitivity analysis in electrophysiological models using multivariable regression. Biophys J. 2009;96: 1264–74. doi:10.1016/j.bpj.2008.10.056

33. Sarkar AX, Sobie EA. Regression Analysis for Constraining Free Parameters in Electrophysiological Models of Cardiac Cells. 2010;6. doi:10.1371/journal.pcbi.1000914

34. Nowotny T, Selverston AI. Models Wagging the Dog : Are Circuits Constructed with Disparate Parameters? 2007;2003: 1985–2003.

35. Weaver CM, Wearne SL. The role of action potential shape and parameter constraints in optimization of compartment models. Neurocomputing. 8th, Feb, 20th ed. 2006;69: 1053–1057. doi:doi:10.1016/j.neucom.2005.12.044 <10.1016/j.neucom.2005.12.044>

36. Brookings T, Goeritz ML, Marder E. Automatic parameter estimation of multicompartmental neuron models via minimization of trace error with control adjustment. J Neurophysiol. 2014;112: 2332–48. doi:10.1152/jn.00007.2014

37. Jȩdrzejewski-Szmek Z, Abrahao KP, Jȩdrzejewska-Szmek J, Lovinger DM, Blackwell KT. Parameter Optimization Using Covariance Matrix Adaptation—Evolutionary Strategy (CMA-ES), an Approach to Investigate Differences in Channel Properties Between Neuron Subtypes. Front Neuroinform. 2018. doi:10.3389/fninf.2018.00047

38. Stein RB, Gossen ER, Jones KE. Neuronal variability: noise or part of the signal? Nat Rev Neurosci. 2005;6: 389–397. 10.1038/nrn1668

39. Jedlicka P, Bird AD, Cuntz H. Pareto optimality, economy–effectiveness trade-offs and ion channel degeneracy: improving population modelling for single neurons. Open Biol. 2022;12. doi:10.1098/rsob.220073

40. Marom S, Marder E. A biophysical perspective on the resilience of neuronal excitability across timescales. Nat Rev Neurosci. 2023;24: 640–652. doi:10.1038/s41583-023-00730-9

41. Zang Y, Marder E. Neuronal morphology enhances robustness to perturbations of channel densities. Proceedings of the National Academy of Sciences. 2023;120. doi:10.1073/pnas.2219049120

42. Hudson A, Prinz A. Conductance ratios and cellular identity. PLoS Comput Biol. 2010;6. doi:10.1371/journal.pcbi.1000838

43. Marder E, Prinz AA. Modeling stability in neuron and network function: the role of activity in homeostasis. Bioessays. 2002;24: 1145–1154. 10.1002/bies.10185

44. Prinz A, Billimoria CP, Marder E. Alternative to Hand-Tuning Conductance-Based Models. Elife. 2014;3.

45. Prinz A a, Bucher D, Marder E. Similar network activity from disparate circuit parameters. Nat Neurosci. 2004;7: 1345–1352. doi:10.1038/nn1352

46. Gunay C, Edgerton JR, Li S, Sangrey T, Prinz AA, Jaeger D. Database analysis of simulated and recorded electrophysiological datasets with PANDORA’s toolbox. Neuroinformatics. 2009/05/29. 2009;7: 93–111. doi:10.1007/s12021-009-9048-z

47. Goaillard J-M, Marder E. Ion Channel Degeneracy, Variability, and Covariation in Neuron and Circuit Resilience. Annu Rev Neurosci. 2021;44: 335–357. doi:10.1146/annurev-neuro-092920-121538

48. Destexhe A, Huguenard J. Which formalism to use for modeling voltage-dependent conductances? In: DeSchutter E, editor. Computational Neuroscience: Realistic Modeling for Experimentalists. Boca Raton, FL: CRC Press; 2000. pp. 129–157.

49. Bower JM, Po-bah H, Beeman D. Looking for Newton: Realistic Modeling in Modern Biology. 2005;1: 1–6. doi:urn:nbn:de:0009-3-2177

50. Marder E, Taylor AL. Multiple models to capture the variability in biological neurons and networks. Nat Neurosci. 2011;14: 133–8. doi:10.1038/nn.2735

51. Druckmann S, Berger T, Hill S. Evaluating automated parameter constraining procedures of neuron models by experimental and surrogate data. Biol Cybern. 2008;99: 371–379. doi:10.1007/s00422-008-0269-2

52. Druckmann S, Hill S, Schürmann F, Markram H, Segev I. A hierarchical structure of cortical interneuron electrical diversity revealed by automated statistical analysis. Cereb Cortex. 2013;23: 2994–3006. doi:10.1093/cercor/bhs290

53. Iavarone E, Yi J, Shi Y, Zandt BJ, O’reilly C, Van Geit W, et al. Experimentally-constrained biophysical models of tonic and burst firing modes in thalamocortical neurons. PLoS Comput Biol. 2019;15. doi:10.1371/journal.pcbi.1006753

54. Arnaudon A, Reva M, Zbili M, Markram H, Van Geit W, Kanari L. Controlling morpho-electrophysiological variability of neurons with detailed biophysical models. iScience. 2023;26. doi:10.1016/j.isci.2023.108222

55. Keren N, Bar-yehuda D, Korngreen A. Experimentally guided modelling of dendritic excitability in rat neocortical pyramidal neurones. J Physiol. 2009;587: 1413–1437. doi:10.1113/jphysiol.2008.167130

56. Korngreen A. Sensitivity analysis of voltage-gated ion channel models. BioRxiv. 2025. doi:10.1101/2025.08.05.668838

57. Bradbury J, Frostig R, Hawkins P, Johnson MJ, Leary C, Maclaurin D, et al. JAX: composable transformations of Python+NumPy programs. 2018. Available: http://github.com/jax-ml/jax

58. Patrick Kidger. On Neural Differential Equations. University of Oxford. 2021.

59. Hamby DM. A review of techniques for parameter sensitivity analysis of environmental models. Environ Monit Assess. 1994;32. doi:10.1007/BF00547132

60. Saltelli A, Ratto M, Andres T, Campolongo F, Cariboni J, Gatelli D, et al. Global sensitivity analysis: The primer. Global Sensitivity Analysis: The Primer. John Wiley & Sons; 2008. doi:10.1002/9780470725184

61. Zi Z. Sensitivity analysis approaches applied to systems biology models. IET Syst Biol. 2011;5. doi:10.1049/iet-syb.2011.0015

62. Sobol IM. Global sensitivity indices for nonlinear mathematical models and their Monte Carlo estimates. Math Comput Simul. 2001;55. doi:10.1016/S0378-4754(00)00270-6

63. Glen G, Isaacs K. Estimating Sobol sensitivity indices using correlations. Environmental Modelling and Software. 2012;37. doi:10.1016/j.envsoft.2012.03.014

64. Herman J, Usher W. SALib: An open-source Python library for Sensitivity Analysis. The Journal of Open Source Software. 2017;2. doi:10.21105/joss.00097

65. Otis TS, Gilly WF. Jet-propelled escape in the squid Loligo opalescens: Concerted control by giant and non-giant motor axon pathways. Proc Natl Acad Sci U S A. 1990;87. doi:10.1073/pnas.87.8.2911

66. Young JZ. The Functioning of the Giant Nerve Fibres of the Squid. Journal of Experimental Biology. 1938;15. doi:10.1242/jeb.15.2.170

67. Pumphrey RJ, Young JZ. The Rates of Conduction of Nerve Fibres of Various Diameters in Cephalopods. Journal of Experimental Biology. 1938;15. doi:10.1242/jeb.15.4.453

68. Clay JR. Excitability of the squid giant axon revisited. J Neurophysiol. 1998;80. doi:10.1152/jn.1998.80.2.903

69. Rosenthal JJC, Bezanilla F. Seasonal variation in conduction velocity of action potentials in squid giant axons. Biological Bulletin. 2000;199. doi:10.2307/1542873

70. Vanier MC, Bower JM. A comparative survey of automated parameter-search methods for compartmental neural models. J Comput Neurosci. 1999;7: 149–171. 10.1023/A:1008972005316

71. Druckmann S, Banitt Y, Gidon A, Schürmann F, Markram H, Segev I. A novel multiple objective optimization framework for constraining conductance-based neuron models by experimental data. Front Neurosci. 2007;1. 10.3389/neuro.01.1.1.001.2007

72. Tasaki I, Watanabe A, Takenaka T. Resting and action potential of intracellularly perfused squid giant axon. Proc Natl Acad Sci U S A. 1962;48. doi:10.1073/pnas.48.7.1177

73. Clay JR. Axonal excitability revisited. Progress in Biophysics and Molecular Biology. 2005. doi:10.1016/j.pbiomolbio.2003.12.004

74. Prescott SA, De Koninck Y, Sejnowski TJ. Biophysical basis for three distinct dynamical mechanisms of action potential initiation. PLoS Comput Biol. 2008;4. doi:10.1371/journal.pcbi.1000198

75. Kim M, McKinnon D, MacCarthy T, Rosati B, McKinnon D. Regulatory evolution and voltage-gated ion channel expression in squid axon: Selection-mutation balance and fitness cliffs. PLoS One. 2015;10. doi:10.1371/journal.pone.0120785

