## Supplemental figures 1 & 2 for "Neuronal excitability and parameter variability in the Hodgkin-Huxley model"

Supplementary information:

Supplementary Figure 1: Distribution of parameter values obtained in the Bootstrap analysis of the Hodgkin-Huxley rate constants. The name of each fitted parameter is indicated below each subplot and is defined in Table 1.

Supplementary Figure 2: Distribution of the four firing subcategories obtained when the simulation was performed with a population of 300000, 800000, and 1200000 parameter sets. The contribution of each subcategory to the population was identical across these simulations.

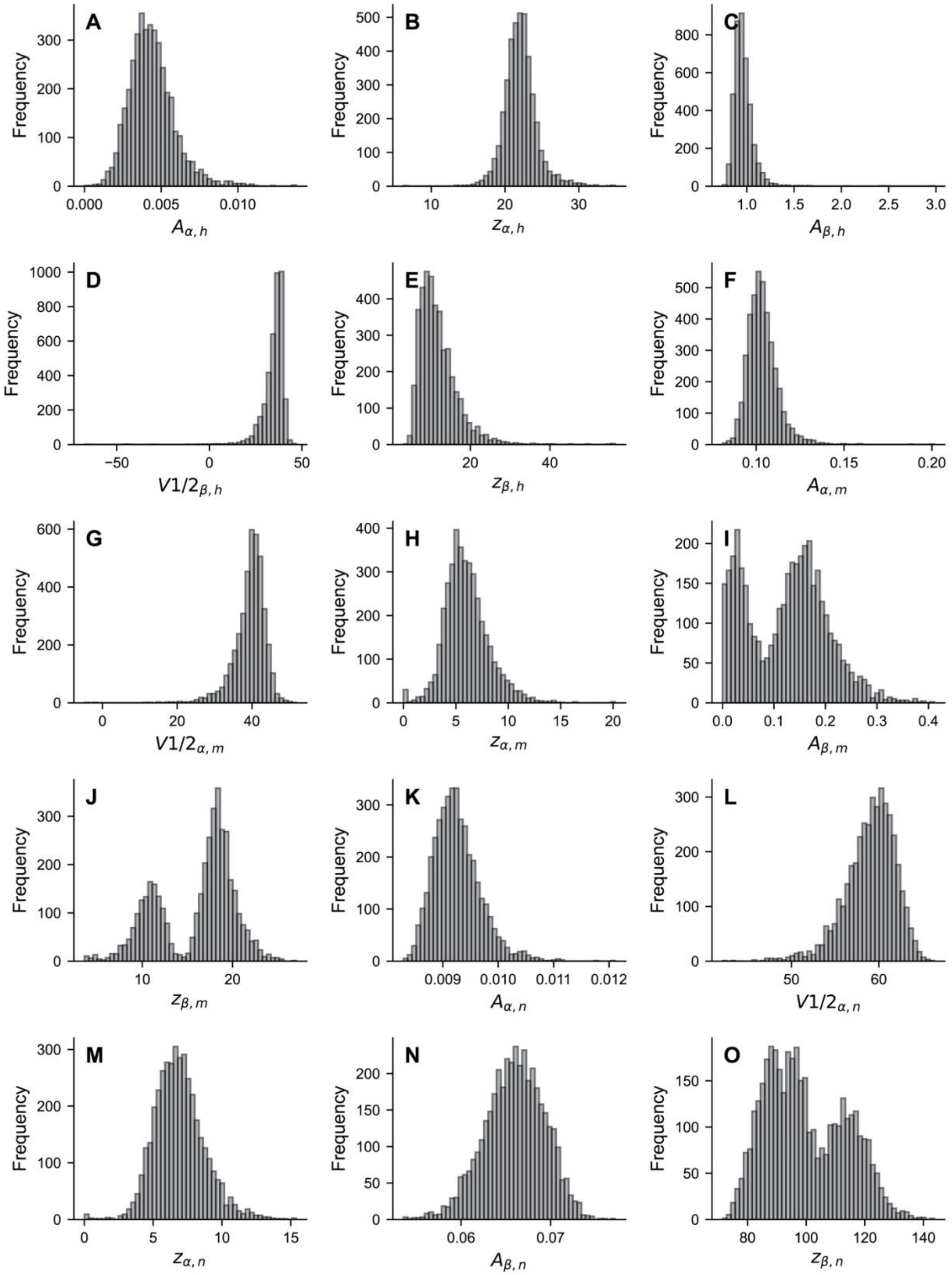


Supplementary Figure 1

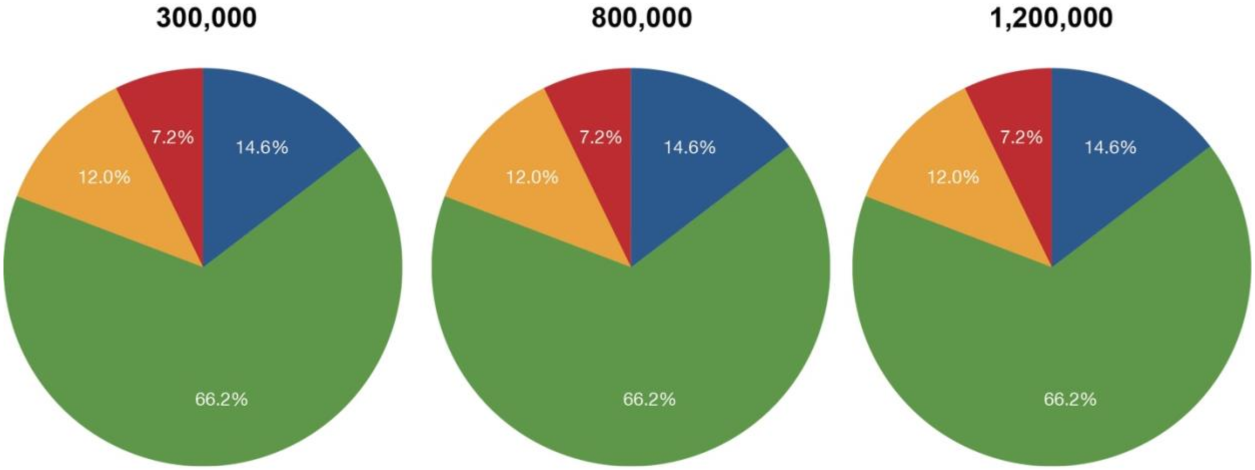


Supplementary Figure 2
